# The tufted ghost crab (*Ocypode cursor*) feeding on stranded loggerhead turtles (*Caretta caretta*) in the southeastern Mediterranean

**DOI:** 10.1101/2025.09.06.674680

**Authors:** Bella S. Galil, Omri Bronstein, Menachem Goren

## Abstract

The tufted ghost crab *Ocypode cursor* is a widespread and ecologically significant inhabitant of Mediterranean sandy beaches, including those of Israel, where it is a characteristic component of the coastal ecosystem. An opportunistic omnivore, *O. cursor* feeds on a broad range of marine and terrestrial resources, including carrion, and has been reported elsewhere preying on sea turtle eggs and hatchlings and scavenging large carcasses. However, feeding on stranded adult sea turtles, particularly loggerhead turtles (*Caretta caretta*), have not previously been documented in the Mediterranean Sea. During a survey assessing ghost crab populations at Zikim Dunes Nature Reserve (southeastern Mediterranean, Israel) in August 2021, we observed five recently stranded *C. caretta* carcasses encircled by freshly dug *O. cursor* burrows, located inland from the high tide line. Burrows surrounding the carcasses were predominantly large-sized, suggesting relocation of adult crabs toward carrion situated landward of their typical supratidal zone. Our findings highlight the opportunistic scavenging behavior of *O. cursor* and underscore their ecological flexibility in exploiting substantial food resources beyond their usual distribution band. These novel observations contribute to understanding ghost crab trophic ecology and their potential interactions with vulnerable marine species in a changing coastal environment.

## 2. Introduction

Loggerhead sea turtles, *Caretta caretta* (Linnaeus, 1758), are categorized as ‘vulnerable’ worldwide (Casale and Tucker, 2017). However, as the Mediterranean subpopulation of the species has been steadily increasing due to successful conservation, its status was altered to ‘Least concern’ (Casale, 2015). Following reports concerning the decimation of the local population, Israel’s Nature and Parks Authority initiated conservation measures in the 1990s consisting of nesting surveys and relocation campaigns (Sella, 1982; Kuller, 1999; Levy and Rybak, 2022). The nesting season occurs from mid-May to the beginning of August, peaking between June and mid-July (Levy, 2005). In 2021, an estimated 170 brown sea turtles and 13 green sea turtles produced 449 nests along the Israeli Mediterranean coast (Levy and Rybak, 2022).

The main threats to the Israeli population of *C. caretta* are fishery bycatch, nesting habitat degradation due to coastal development and recreational use, and eggs and hatchlings predation by crows, dogs and foxes (Leader et al., 2022). Predation by the tufted ghost crab, *Ocypode cursor* (Linnaeus, 1758), on the eggs and hatchlings, though recorded elsewhere as drivers of high mortality rates (Smith et al., 1996; Strachan et al., 1999, Marco et al., 2015), has not been reported from the Israeli coast. Similarly, scavenging by *O. cursor* on adult loggerhead turtles has never been reported from the Mediterranean Sea.

Fecundity in brachyuran crabs is strongly correlated with female body size, which serves as the primary determinant of reproductive output. For example, Hines (1991) found that despite interspecific differences in egg size and body mass, the allometric relationship between body size and fecundity was consistent across nine species of cancerid crabs in the North Pacific and Atlantic, reflecting an evolutionary pattern in which larger body size facilitates increased reproductive investment. Spatial allocation by larger, sexually mature individuals, toward areas with higher availability of opportunistic food sources, such as stranded carrion, may thus enhance reproductive potential by improving energy intake and supporting the metabolic demands of egg production. Consequently, opportunistic food sources my drive adult crabs to relocate beyond their natural distribution range.

During the summer of 2021, while assessing the effects of recreational disturbance on *O. cursor* populations in sandy shore nature reserves (Galil et al., 2024), we noted 5 carcasses of recently deceased loggerhead sea turtles surrounded by freshly dug crab burrows, suggesting opportunistic scavenging by *O. cursor*, with adult crabs relocating and aggregating around substantial carrion resources beyond their typical supratidal zone.

## 3. Material and methods

The sandy shore of Zikim Dunes nature reserve (31.6297 N, 34.5151 E – 31.6148 N, 34.5053 E), established in 2005, is the southernmost site along the Israeli coastline where *C. caretta* nesting were reported. The shore has no recreational facilities and thus not subject to substantial recreational activities even during peak coastal recreation season during the summer (June–September). The abundance and population size structure of *O. cursor* were surveyed using nondestructive methods during August 2021. A 15 m wide transect parallel to the water edge was surveyed for the presence of burrows and measurement of burrow opening diameter (BOD) following the protocol of Galil et al. (2024). BOD is widely accepted as a non-intrusive and environmentally benign proxy for estimating resident crab size, given its established correlation with crab body size (Shuchman and Warburg, 1978; Strachan et al., 1999; Türeli et al., 2009). Each burrow is typically occupied by a single crab (Lewinsohn, 1964). Burrow openings were examined for signs of freshly dug sand and/or marks left by crabs’ dactyls to determine whether the burrow has been in recent use. BOD was measured to the nearest mm.

## 4. Results

Five carcasses of *C. caretta* sea turtles in early stages of decomposition (bloody secretions were observed near two individuals) were found on August 26, 2021, on the shore of Zikim Dunes nature reserve, encircled with freshly dug crab burrows (Fig. 1). One carcass was found next to the high tide line (HTL) (Table 1, specimen 1), others were strewn landwards, at distances approximately 20-25 m beyond the HTL. Those carcasses were surrounded with between 8 and 18 burrows with BOD ranging from 20-60 mm) (Table 1, specimens 2-5).

**Table 1.** Number and burrows opening diameter (BOD) of *Ocypode cursor* encircling *Caretta caretta* carcasses in Zikim Dunes nature reserve, Israel, 26^th^ August 2021.

|  | Specimen |  |  |  |  |
| --- | --- | --- | --- | --- | --- |
| *Caretta caretta* | 1 | 2 | 3 | 4 | 5 |
| Number of encircling burrows | 2 | 15 | 17 | 8 | 18 |
| Minimum/Maximum BOD (mm) | 11/30 | 20/50 | 25/60 | 12/47 | 30/50 |
| Average BOD (mm) |  | 37.27 | 42.59 | 35.50 | 42.39 |
| S.D. |  | 10.02 | 8.44 | 11.49 | 6.70 |

**Figure 1.**
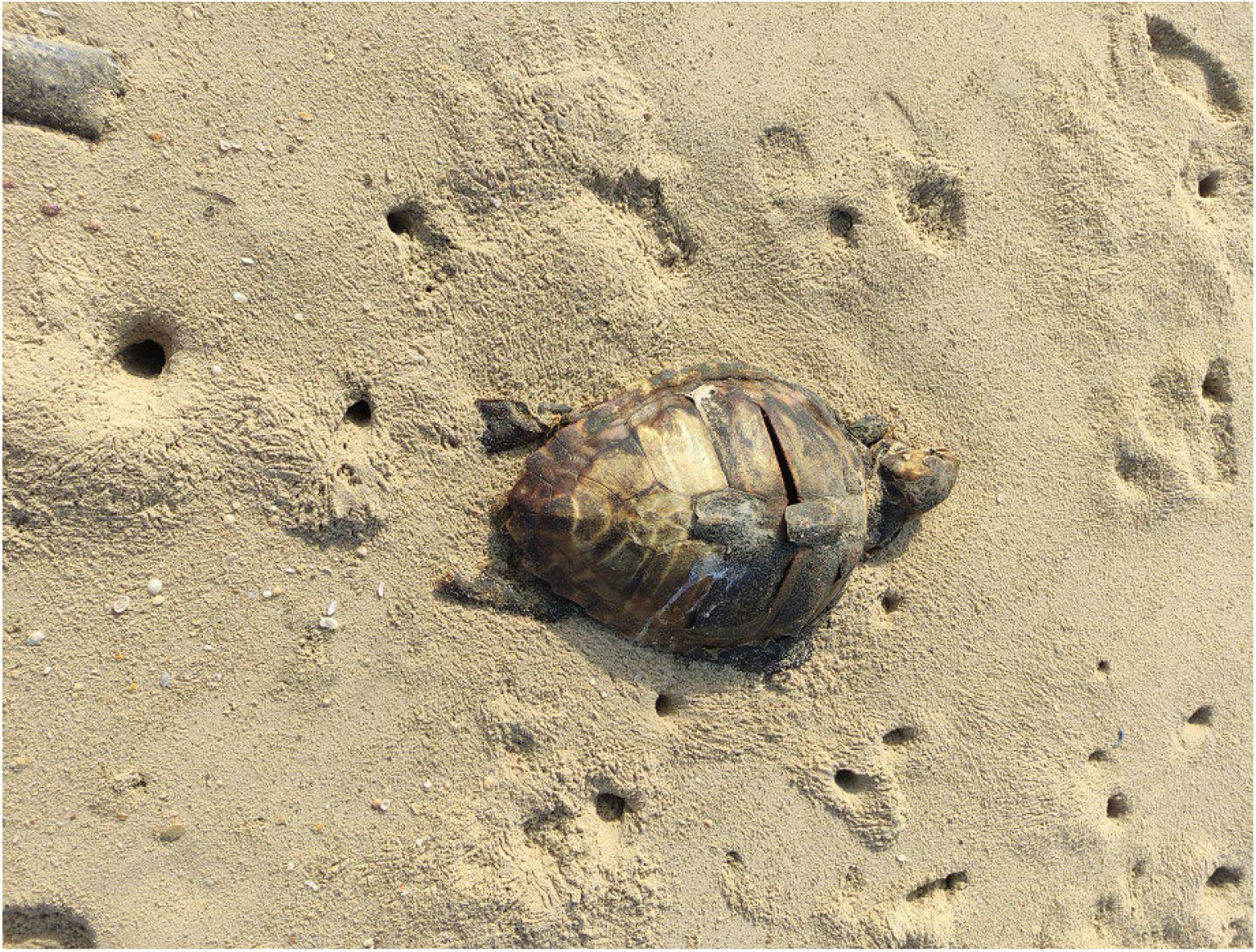
*Caretta caretta* (Linnaeus, 1758) carcass surrounded by active burrows of *Ocypode cursor* (Linnaeus, 1758), 26 August 2021, Zikim Dunes nature reserve, along the southern Mediterranean coast of Israel. Photo K. Gayer.

At all four inland carcasses, BOD of the carcasses-scavenging crabs were larger than those documented that same day at the adjacent Nitzanim nature reserve (Galil et al. 2024). Larger individuals equipped with larger, more robust claws, are capable of breaching the tough integument of turtle carcasses, thereby gaining access to a nutrient-rich food source.

## 5. Discussion

*Ocypode cursor* burrows along the microtidal Mediterranean coast of Israel are clustered along a narrow band above the tide line, with burrows only rarely observed along the landward edge – approximately 20 m away from the swash zone. The proportion of large sized burrows (BOD > 41 mm) increases landwards whereas burrows of juvenile crabs (BOD < 20 mm) decrease with growing distance from the waterline (Galil et al., 2024). Similarly, Schuchman and Warburg (1978) reported that ‘The smaller burrows … were mostly found closer to the sea; the [larger-sized] group was found more landwards’. Surveys of *O. cursor* conducted in Cyprus and Turkey concur, showing most small burrows closest to the tide line, and largest burrows found farthest (Barakalı et al., 2020; Strachan et al., 1999; Türeli et al., 2009, 2014).

With respect to *C. caretta*, nest placement is typically positioned at a significant distance inland from the high tide line (Hays and Speakman, 1993), beyond the band of *O. cursor* burrows. This ecological zonation may be the reason that *O. cursor* target hatchlings traversing the beach en route to the sea and only occasionally prey on nests located landwards of their zone, thus affecting only a very small proportion of nests (Erk’akan, 1993; Smith et al., 1996; Strachan et al., 1999; Türkozan et al., 2003). Ghost crabs are opportunistic predators and facultative scavengers with an omnivorous diet: *O. cursor* preys on both marine (crustaceans and molluscs) and terrestrial organisms, macroalgae, as well as leftover human food and carrion (Strachan et al., 1999; Chartosia et al., 2010; Avenant et al., 2024; Marchessaux et al., 2024). Strachan et al. (1999: 56) observed *O. cursor* feeding upon a cow carcass located “…relatively high up the beach, above the normal burrow zone. Yet, surrounding the carcass were crab burrows, all with openings > 35 mm in diameter.” The only *O. cursor* burrow clusters encountered beyond the supratidal band during our survey were those encircling the four *C. caretta* carcasses, and the greater majority were large sized adults (BOD > 41 mm), larger than those encountered in August 2021 in the nearby Nitzanim nature reserve (Galil et al., 2024, Table 3). It appears that large-sized individuals on encountering a sizable carcass would relocate and resettle in its vicinity, even if it is situated further inshore than their habitual/typical location.

## Acknowledgements

This note is dedicated to Yigal Sella, Zeev Kuller and Yaniv Levy in appreciation for their decades-long efforts to restore the local populations of marine turtles. Special thanks to Kfir Gayer for the ‘beachcombing’ summer of 2021.

